# High-throughput cell-based assays for the preclinical development of DsbA inhibitors as antivirulence therapeutics

**DOI:** 10.1101/2020.03.03.975938

**Authors:** Anthony D. Verderosa, Rabeb Dhouib, Yaoqin Hong, Begoña Heras, Makrina Totsika

## Abstract

Antibiotics are failing fast, and the development pipeline is alarmingly dry. New drug research and development is being urged by world health officials, with new antibacterials against multidrug-resistant Gram-negative pathogens as the highest priority. Antivirulence drugs, which are inhibitors of bacterial pathogenicity factors, are a class of promising antibacterials, however, their development is often stifled by lack of standardised preclinical testing akin to what guides antibiotic development. The lack of established target-specific microbiological assays amenable to high-throughput, often means that cell-based testing of virulence inhibitors is absent from the discovery (hit-to-lead) phase, only to be employed at later-stages of lead optimization. Here, we address this by establishing a pipeline of bacterial cell-based assays developed for the identification and early preclinical evaluation of DsbA inhibitors. Inhibitors of DsbA block bacterial oxidative protein folding and were previously identified by biophysical and biochemical assays. Here we use existing *Escherichia coli* DsbA inhibitors and uropathogenic *E. coli* (UPEC) as a model pathogen, to demonstrate that a combination of a cell-based AssT sulfotransferase assay and the UPEC motility assay, modified for a higher throughput format, can provide a robust and target-specific platform for the evaluation of DsbA inhibitors. Our pipeline could also be used in fragment and compound screening for the identification of new DsbA inhibitor classes or hits with a broad spectrum of activity. In conclusion, the establishment of accurate, high-throughput microbiological assays for antivirulence drug identification and early preclinical development, is a significant first step towards their translation into effective therapeutics.

**Importance:** The safety net of last resort antibiotics is quickly vanishing as bacteria become increasingly resistant to most available drugs. If no action is taken, we will likely enter a post-antibiotic era, where common infections and minor injuries are once again lethal. The paucity in new antibiotic discovery of the past decades has compounded the problem of increasing antibiotic resistance, to the point that it now constitutes a global health crisis that demands global action. There is currently an urgent need for new antibacterial drugs with new targets and modes of action. To address this, research and development efforts into antivirulence drugs, such as DsbA inhibitors, have been ramping up globally. However, the development of microbiological assays as tools for effectively identifying and evaluating antivirulence drugs is lagging behind. Here, we present a high-throughput cell-based screening and evaluation pipeline, which could significantly advance development of DsbA inhibitor as antivirulence therapeutics.

## Introduction

In 2014 the World Health Organisation (WHO) released a statement declaring antimicrobial resistance (AMR) as a public health priority that demands decisive global action (1). Although WHO’s statement has increased AMR awareness, at the time of writing over half a decade has passed, and little progress has been made in developing effective solutions (2, 3); meanwhile, AMR rates continue to rise. The current AMR crisis demands the urgent development of effective strategies to tackle bacterial infections. One actively researched strategy is the development of antivirulence therapeutics, which have recently been gaining momentum as effective antibacterials that can circumvent the mechanisms of antibiotic resistance (4). Antivirulence drugs target bacterial virulence factors and are designed to disarm pathogens, unlike conventional antibiotics which either kill or inhibit bacterial growth (5, 6). Targeting virulence factors can attenuate a pathogen’s ability to cause infection and render bacteria susceptible to the host’s defence systems (7). Consequently, virulence factors present a plethora of attractive targets for the development of new therapeutics.

Although several antivirulence drugs are currently under various stages of development, (e.g. toxin, adhesin, enzyme, secretion and quorum sensing inhibitors (6, 8, 9)) the potential of any antivirulence drug candidate for further clinical development relies on having established robust assays for evaluating their efficacy *in vitro* and *in vivo* (10). While the development of antibiotics over the past several decades, has benefited from standardised and comprehensive preclinical and clinical evaluation methods, the field of antivirulence drugs has had minimal guidelines for consistent testing, with only a few general guidelines reported for some types of inhibitors, e.g. for quorum sensing (10, 11). In addition, antivirulence inhibitor screening campaigns often utilise biophysical and/or biochemical assays (when the target is known), which do not allow early evaluation of inhibitor effects on bacterial cells (12), or on cell-based virulence assays (target agnostic), which might be prone to bias by reporting non-specific inhibitor effects (11). Here we develop a pipeline of robust cell-based assays for the *in vivo* evaluation of inhibitors against the DsbA antivirulence target.

In Gram-negative pathogens, the biogenesis and function of many virulence factors are intrinsically linked to the redox enzyme pair of DsbA and DsbB (13–16). DsbA is a periplasmic oxidoreductase which catalytically introduces disulfide bonds into secreted and outer membrane proteins (17), while its inner membrane partner DsbB reoxidises DsbA (18, 19). Intramolecular disulfide bonds are often essential for the native folding and subsequent function of multiple secreted or surface proteins, including fimbriae, flagellar motor, secretion systems, and secreted toxins (13, 16). Given that many of these proteins are *bona fide* virulence factors or form integral components of machinery for virulence factor assembly, this makes DsbA and DsbB ideal targets for the development of antivirulence drugs (13, 16, 20). Recently, several classes of small molecule inhibitors of DsbA, as well as inhibitors of its cognate DsbB, have been reported, primarily through screening campaigns involving biophysical and/or biochemical assays (12, 21–25). Any *in vivo* assessment of promising hits was typically conducted as part of subsequent testing, often at a stage where significant efforts into the chemical elaboration of initial hits had already taken place. Incorporation of cell-based testing at an earlier stage of inhibitor screening, as conducted for DsbB and its homologue VKOR (24), could be used to complement early hit selection by biophysical/biochemical approaches and likely save time and money, by informing which hits should be prioritised and what properties should be optimised (e.g. solubility, cell permeability, toxicity etc.).

For monitoring DsbA function *in vivo*, the bacterial motility assay on soft agar has been most commonly used (26–28) and more recently this method was applied to DsbA inhibitor testing *in vivo* (12, 29). In many pathogens, such as uropathogenic *Escherichia coli* (UPEC), and *Salmonella enterica* serovar Typhimurium (*S.* Typhimurium), motility requires the production of functional flagella, with DsbA playing a central role in the biogenesis of these surface appendages (26, 30–33). The standard bacterial motility assay format (performed in Petri dishes) is however relatively low-throughput and requires large inhibitor quantities and manual data collection (29), thus, limiting its utility for high-throughput inhibitor screening and testing. A second method recently utilised for DsbA inhibitor testing monitors the enzymatic activity of AssT (29), an arylsulfate sulfotranserase encoded by several pathogens (e. g. UPEC, *S.* Typhimurium, *Klebsiella* (27, 28, 34, 35)), which is proposed to play a role in the intracellular detoxification of phenolic substances (36–38). AssT is a native substrate for DsbA and its homologue DsbL (39) as it requires the formation of a disulfide bond for its correct function (40). Consequently, AssT sulfotransferase activity can be used to measure DsbA activity *in vivo*, and can be monitored either in solution (39) or using an agar-based assay (27). Although very informative, previously used AssT assays have not been amenable to high-throughput inhibitor screening and testing. Here, we present a comprehensive pipeline of cell-based assays that provide an accurate and high throughput platform for the identification of DsbA inhibitors and their subsequent development, from hits to leads, and from lead optimisation to early preclinical candidate validation.

## Results

### *E. coli* motility can be accurately tracked spectrophotometrically

As bacterial motility assays are the most common method of assessing DsbA inhibitor efficacy, we first sought to develop a motility assay which would circumvent the limitations of the standard assay format and provide a platform which could be utilised earlier on in the drug development pipeline. We adapted a microtiter plate-based assay which had previously been reported for screening antimicrobial compounds using bacterial motility (41). We first confirmed that bacterial swimming motility could be accurately monitored spectrophotometrically. As bacteria radially migrated thought the soft agar, a zone of motility corresponding to an increase in absorbance at 600 nm was observed, and a motility curve could be generated over time (Figure 1). Using this method, the start, end, and motility rate (slope) of the tested *E. coli* strain (JCB816) could be accurately measured under a set of specific culture conditions.

**Figure 1.**
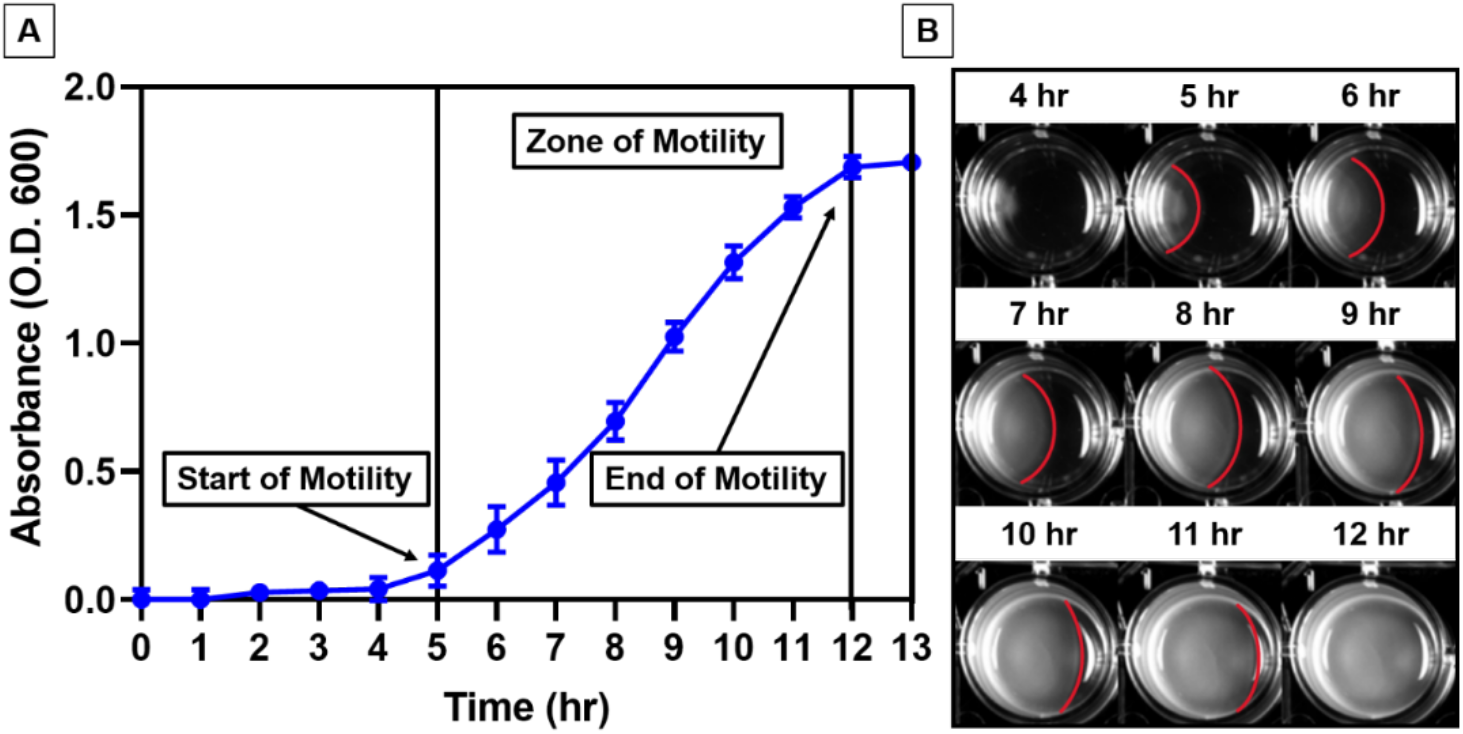
Absorbance-based monitoring of E. coli motility. (A) Motility curve of E. coli JCB816 monitored spectrophotometrically during incubation on soft LB agar at 37°C over 15 hours, as detailed in methods. Data points represent hourly mean absorbance values ± SD from 3 independent culture replicates. (B) Digital images tracking the swimming motility of E. coli JCB816 on soft LB agar in a 24-well plate. E. coli was inoculated at the left edge of each well, and by 12 hours incubation at 37°C the zone of motility (boundary marked in red) had reached the opposite edge of the well.

### An absorbance-based bacterial motility assay optimised for DsbA inhibitor evaluation

To demonstrate the value of a plate reader-based motility assay in assessing antivirulence DsbA inhibitor hits, we generated motility curves for UPEC strain CFT073 in the presence and absence of phenylthiophene inhibitor F1 (Figure 2C), which we have previously shown to inhibit DsbA in CFT073 using the traditional petri-dish motility assay (29). The motility of CFT073 in soft agar containing the DsbA inhibitor F1 at a concentration gradient (1-0.1 mM) was reduced compared to the vehicle control in a dose-dependent manner, with maximum motility inhibition observed at 1 mM F1 (97% compared to DMSO control at 10 hours post-inoculation) (Figure 2A). Analysing longitudinal motility data revealed that both the start time and the rate of motility are directly related to F1 inhibitor concentration, with higher concentrations resulting in longer motility start times and slower motility rates (Table 1) - effects that were previously evident with the conventional motility assay methodology (26). Furthermore, the high reproducibility of our assay allowed for even small changes in motility rate to be robustly detected (*P* <0.0001, one-way ANOVA test) between different F1 treatment groups (Table 1). Using the calculated rate of motility for each F1 concentration, the F1 dose-response curve was generated (Figure 2B) with the half maximal inhibitory concentration (IC_50_) of F1 calculated at the 0.35-0.47 mM range. IC_50_ values in the mM range are indicative of modest affinity inhibitor hits that represent good candidates for synthetic optimisation. Taken together, our absorbance-based motility assay proved to be of value in generating accurate and highly reproducible motility curve data that could be used to identify and characterise early hits from DsbA inhibitor screening campaigns, such as inhibitor F1 (12). Moreover, this motility assay format (24-well plate) required almost 29-fold less inhibitor than the standard petri-dish assay (0.14 mg/well versus 4 mg/petri-dish) and used an automated data collection pipeline that markedly reduced assay hands-on time. Optimising the assay for a 24- or 48- well plate format also increases the applicability of the assay for medium-throughput inhibitor screens, however, this assay cannot be optimally adapted for high-throughput screening (e.g. 96 or 384-well plate format), which would be typically employed in early screening campaigns using large fragment/compound libraries.

**Figure 2.**
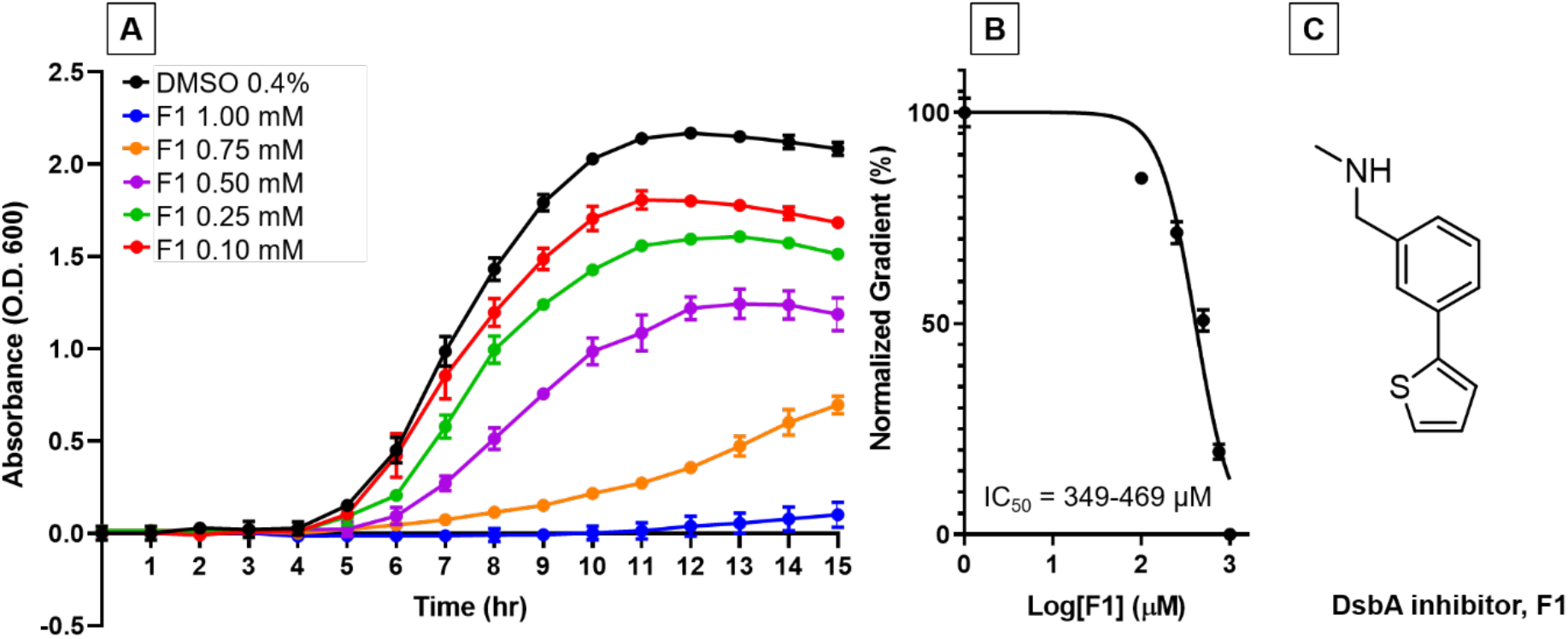
Absorbance-based UPEC motility in varying concentrations of DsbA inhibitor F1. (A) Motility curves and (B) motility dose-response curve of UPEC CFT073 on LB agar (0.25%) containing DsbA inhibitor F1 (1-0.1 mM) or 0.4% DMSO (vehicle control), generated as detailed in methods. (C) chemical structure of DsbA inhibitor F1. Data points represent the mean ± SD of 3 biological replicates.

**Table 1.** Motility curve parameters for UPEC CFT073 in the presence of varying concentrations of DsbA inhibitor F1.

| F1 (mM) | Start of motility (hr) | End of motility (hr) | Rate of motility (slope)<sup>a</sup> | Inhibition of motility (%)<sup>b</sup> |
| --- | --- | --- | --- | --- |
| 1 | 11 | n/d | 0.02±0.007 | 97 |
| 0.75 | 7 | n/d | 0.09±0.013 | 89 |
| 0.5 | 6 | 12 | 0.21±0.02 | 51 |
| 0.25 | 5 | 11 | 0.28±0.02 | 30 |
| 0.1 | 5 | 11 | 0.33±0.01 | 17 |
| 0 | 5 | 11 | 0.38±0.03 | n/a |

<sup>a</sup>motility rates shown as mean slope value ± S.D. from four biological replicates. Group means were compared using the one-way ANOVA test ( $P < 0.0001$ ).
<sup>b</sup>compared to vehicle control (DMSO 0.4%) and determined using data from the 10 -hour time point
n/d = not determined.

### Establishing a high throughout cell-based enzymatic assay for DsbA inhibitor screening and development

Enzymatic assays are better suited to high-throughput inhibitor screening campaigns. Thus, we sought to develop a cell-based assay for monitoring the activity of the AssT enzyme, which is a native DsbA substrate in UPEC. We first determined if AssT’s sulfotransferase activity could be assayed in solution using live UPEC cells cultured in standard laboratory conditions. The AssT overexpressing strain CFT073/pAssT was cultured overnight in LB (with appropriate selection to maintain the pAssT vector or pSU2718 vector control), and culture aliquots were mixed in a 96-well plate with the aryl sulfate phenolic donor, 4-methylumbelliferyl sulfate (MUS) and the phenol acceptor, phenol (detailed in methods and Figure 6). AssT catalysed the cleavage of the sulfate group from the non-fluorescent substrate MUS to the highly fluorescent product 4-Methylumbelliferone (MU) (Scheme 1) (42). A steady increase in fluorescence was observed over time for strain CFT073/pAssT, but not for the vector control (Figure 3A), confirming production of functional AssT enzyme which catalysed the conversion of MUS and phenol in the UPEC periplasm (where AssT and DsbA localise) *in vivo*. Retaining LB growth medium in the assay reaction, which significantly reduced assay time and labour-intensiveness, did not block or interfere with the fluorescence output of the sulfotransferase reaction (data not shown).

**Scheme 1.**
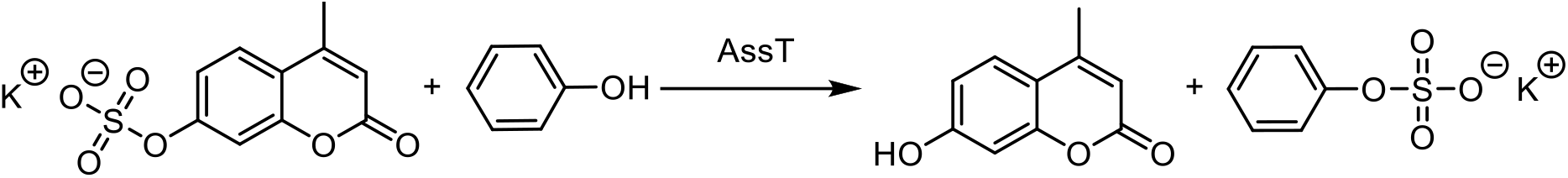
AssT catalysed conversion of MUS to MU.

**Figure 3.**
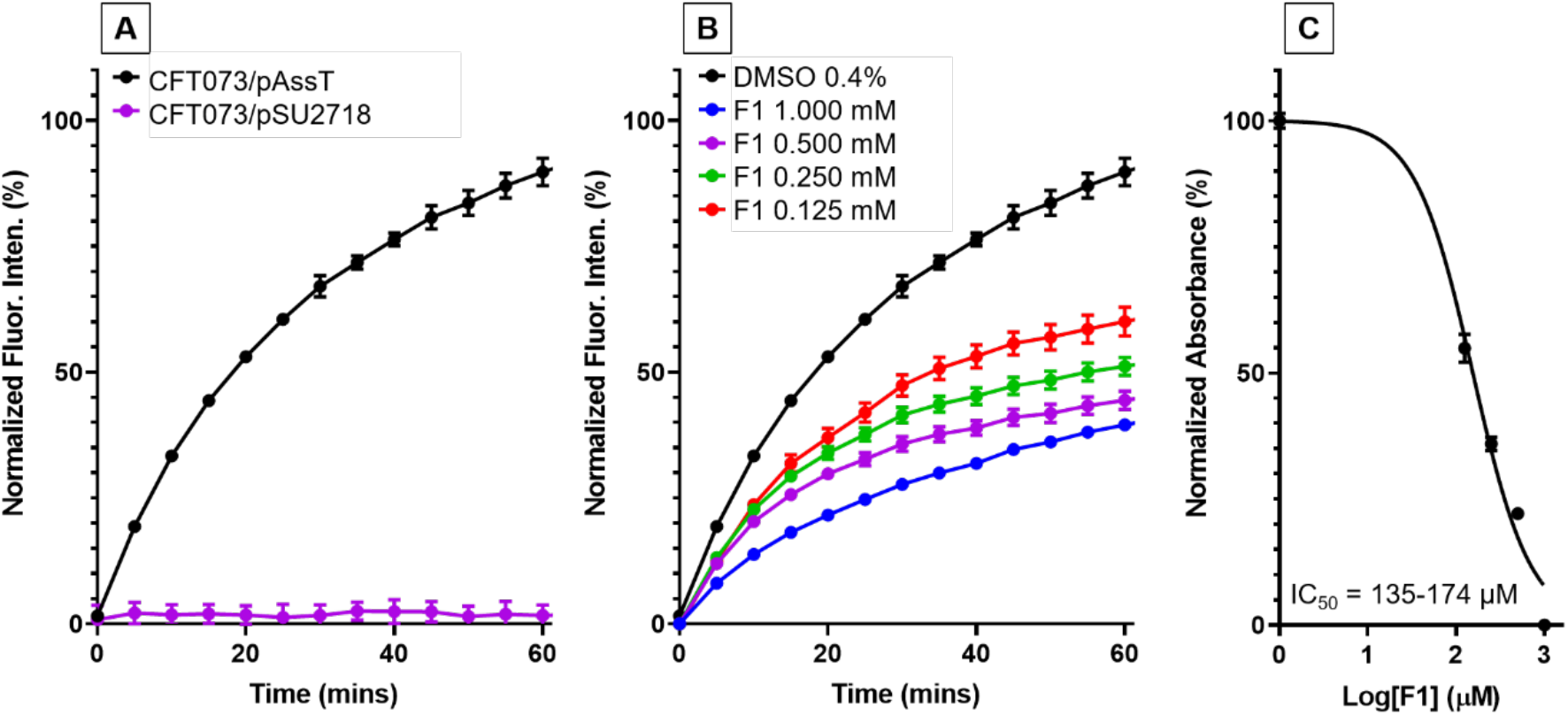
Cell-based AssT activity in UPEC CFT073 in varying concentrations of DsbA inhibitor F1. Sulfotransferase activity of (A) CFT073/pASST and CFT073/pSU2718 (vector control), (B) CFT073/pAssT, and (C) corresponding dose-response curve (calculated at 40-minute time point) cultured in the presence of F1 (1-0.125 mM) or 0.4% DMSO (vehicle control). F1 treated bacterial cultures were mixed with MUS and phenol and immediately monitored spectrofluorometrically as detailed in methods. Data are shown as normalised fluorescence intensity units, with the mean ± SD of 3 biological replicates plotted at each time point.

### The cell-based AssT sulfotransferase assay offers a high-throughput platform for DsbA inhibitor development

We hypothesised that inhibition of DsbA in CFT073/pAssT would result in misfolding of the AssT enzyme and loss of sulfotransferase activity. To examine this hypothesis, we repeated the AssT assay with CFT073/pAssT cells treated with various concentrations of DsbA inhibitor F1 (1-0.125 mM). Sulfotransferase activity was significantly decreased at all tested F1 concentrations (Figure 3B), with lowest fluorescence measured from cells cultured at an F1 concentration of 1 mM (55% reduction compared to vehicle (DMSO) control). Reduction of AssT sulfotransferase activity by F1 was dose-dependent, and from the dose-response curve F1 had an IC_50_ value in the 0.14-0.17 mM range (Figure 3C). This was similar to F1’s IC_50_ value range calculated from motility data, suggesting that the two assays are reporting similar DsbA inhibition of two independent virulence substrates. Taken together, these results demonstrate that DsbA inhibition results in loss of AssT sulfotransferase activity, confirming that our cell-based AssT sulfotransferase assay can be used for indirectly assessing DsbA inhibition in a high-throughput format.

### The cell-based AssT enzyme assay allows target-specific testing of DsbA inhibitor activity

With assay protocol and conditions optimised, we next sought to confirm the specificity of our AssT assay for the DsbA target. To investigate this, we utilised a previously characterised CFT073 mutant lacking DsbA and DsbL (a DsbA homologue encoded by UPEC (28)), which was transformed with a plasmid carrying the AssT enzyme (CFT073Δ*dsbA*Δ*dsbLI*/pAssT). This strain lacking both DsbA homologues had significantly decreased fluorescence compared to the wild-type strain (CFT073/pAssT). *In trans* complementation with DsbA fully restored the mutant’s fluorescence back to wild-type levels (Figure 4A), confirming that in our assay DsbA is required for the production of functional AssT enzyme. In addition, both the control strain CFT073/pAssT and the complemented mutant CFT073Δ*dsbA*Δ*dsbLI*/pAssT/pEcDsbA were equally attenuated for AssT function when treated with 0.5 mM F1 inhibitor (Figure 4A). In contrast, the mutant (CFT073Δ*dsbA*Δ*dsbLI*/pAssT) was unresponsive to F1 treatment, and its fluorescence profile remained unaltered upon treatment with 0.5 mM F1 or with DMSO (Figure 4A). These results confirm that our assay can identify inhibitors that specifically target DsbA, as DsbA is the main factor mediating high levels of functional AssT enzyme.

**Figure 4.**
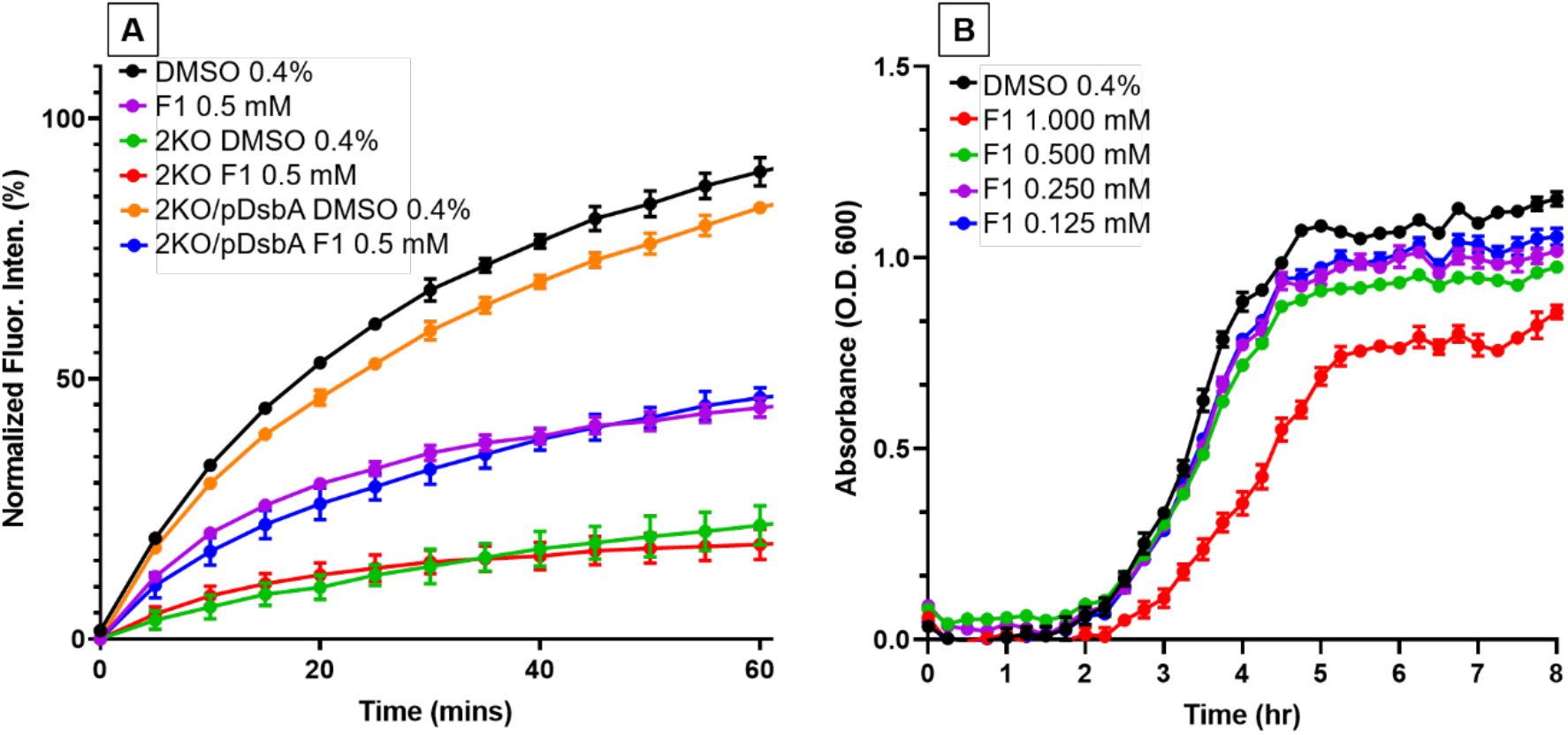
F1 inhibitor effects on UPEC DsbA function and growth. (A) Cell-based sulfotransferase activity of CFT073/pAssT grown in the presence of 0.4% DMSO (black) or 0.5 mM F1 (purple); CFT073ΔdsbAΔdsbLI/pAssT (2KO) grown in the presence of 0.4% DMSO (green) or 0.5 mM F1 (red); and CFT073ΔdsbAΔdsbLI/pAssT/pEcDsbA (2KO/pDsbA) grown in the presence of 0.4% DMSO (orange) or 0.5 mM F1 (blue). (B) Growth curves of CFT073/pAssT cultured in LB medium containing F1 (1-0.125 mM) or 0.4% DMSO (vehicle control) and monitored spectrophotometrically (Optical Density (O.D.) at 600 nm) as detailed in methods. Data are normalised fluorescence intensity units (A) or absorbance at 600 nm (B), with mean ± SD of 3 biological replicates plotted at each time point.

### Adding a growth analysis step prior to assessing sulfotransferase activity in the cell-based AssT assay simultaneously screens for inhibitor effects on DsbA function and bacterial growth

DsbA is not required for UPEC growth in rich media and standard laboratory culture conditions (28). As such, inhibitors specific to DsbA would be predicted to have no effect on UPEC growth under these conditions. On the other hand, large libraries of low affinity compounds, such as those typically used in early inhibitor screens, could contain several compounds with bacterial growth toxicity. In order to incorporate growth testing as part of our high-throughput cell-based AssT assay, UPEC growth was continuously monitored (step 1) during culture for the preparation of live-cell samples for sulfotransferase activity testing (step 2). Testing F1 in the growth analysis step of the sulfotransferase assay, revealed that UPEC growth was slightly reduced in the presence of 1 mM F1, with no growth defects observed at lower F1 concentrations (0.5-0.125 mM) (Figure 4B). In addition to uncovering this small growth defect at high F1 concentration, incorporating the growth step in our assay allowed us to account for any potential reduction in viable cells present in culture samples tested for sulfotransferase activity. Having an accurate O.D. 600 nm reading at the time of culture collection, ensured that all samples tested in the AssT assay could be easily adjusted to contain the same number of live cells, which was confirmed by plating samples for viable CFU (data not shown). These results demonstrate that adding a growth analysis step to the cell-based sulfotransferase enzyme assay allows growth related inhibitor effects to be identified and corrected prior to downstream inhibitor testing.

## Discussion

Antimicrobial drug development typically starts with screening large fragment or compound libraries to identify initial hits and the subsequent chemical elaboration of different hit series. Such screening campaigns represent a big investment, in terms of time and resources, both for the industry and for the academic lab. Success relies heavily on the use of well-established, accurate reporter assays that can identify hits with some degree of target-specificity and are amenable to high-throughput testing of several thousands of compounds at once. For antibiotics, such testing is now considered routine and follows global standards and guidelines (43, 44). For non-traditional antibacterials, however, which are currently being actively explored as viable solutions to the pressing problem of AMR, consistency in drug testing and reporting is far from achieved. For antivirulence drugs in particular, a major challenge lies in standardising preclinical testing for a largely diverse set of targets that potentially mediate multiple different phenotypes in bacterial pathogens. Measuring virulence target inhibition reliably and at large-scale is often difficult when using microbiological assays, so when the target is known, inhibitor screening and early evaluation typically relies on biochemical/biophysical approaches. This is the case for DsbA inhibitors that have been reported to date, with hits from several chemical classes having been identified as part of fragment-based screening campaigns primarily using saturation transfer distance NMR spectroscopy (12, 21–23). Later-stage microbiological evaluation has validated some but not all hits, and in some cases, even chemically elaborated analogues have failed to show activity in cell-based assays (12, 29 and unpublished data). In this study, we have adapted two cell-based assays previously used to monitor DsbA function *in vivo* for accurate and high-throughput testing of DsbA inhibitors. When combined, these assays could support DsbA inhibitor development from hit identification to lead optimisation and preclinical candidate validation.

Flagella-mediated bacterial motility is a useful reporter phenotype for DsbA activity and thus the standard petri-dish soft agar motility assay has been successfully used to evaluate DsbA inhibitors *in vivo* (12, 29). However, the current format of the bacterial motility assay in petridishes has several limiting factors, which are preventing its use in inhibitor screens: (i) a relatively low-throughput capacity, (ii) the requirement of high inhibitor quantities, and (iii) manual data collection by either incremental or single endpoint imaging (27, 29, 31). Our modified plate-reader motility assay utilises the same soft agar methodology, however, instead of relying on incremental images and manual measurements of motility zones, it uses a fully-automated system (plate-reader absorbance measurements) and requires no human intervention throughout the assay period making it less labour-intensive and less prone to bias or human error in motility assessments. In addition, automating the assay ensures conditions (e.g. temperature) are better controlled and can remain constant from start to finish. An important improvement was in that downscaling the assay from a petri-dish to a multi-well plate format, drastically reduced the quantity of inhibitor required (30-fold reduction in 24-well plate and 60-fold reduction in 48-well plate, compared to previous method (29)). While others have demonstrated that agar-based motility assay can be performed in 96-well or even a 384-well plate (41, 45), we found that reducing the well diameter below 11 mm (48-well plate) drastically reduced within assay reproducibility, especially when testing inhibitors (data not shown). For this reason, our absorbance-based motility assay is better suited to inhibitor testing post-hit discovery, leaving the need for developing another cell-based DsbA reporter assay that was amenable to high throughput screening of DsbA inhibitors.

For developing such a high throughput assay, we chose to use a read-out that is a native virulence substrate of DsbA in UPEC. AssT is a large periplasmic enzyme encoded by UPEC and other intestinal bacteria (29) that was reported to be upregulated in the urine of UPEC-infected mice (46, 47), but was not required for colonisation of the murine bladder (29). The gene encoding AssT is found in a tri-cistronic operon with the *dsbL* and *dsbI* genes, which encode an accessory redox protein pair in UPEC with specificity for AssT (39, 48), although the DsbA and DsbB redox pair was also shown to functionally fold AssT (27, 29). The AssT activity assay was previously performed in liquid medium using bacterial cell lysates (39) or on solid medium using whole live cells (27). To evaluate DsbA inhibitors, we have previously utilised the solid medium cell-based method to successfully quantify DsbA inhibition of AssT activity (29). Despite this being an accurate cell-based assay, its petri-dish format presented the same limitations as the standard bacterial motility assay above. The modified AssT sulfotransferase enzyme assay presented here operates on the same principle, yet its application is quite different. The assay is conducted in liquid media using live UPEC cells treated with minimal quantities of DsbA inhibitor, and the activity of AssT is assessed in an automated fashion by monitoring the MUS-phenol sulfotransferase reaction spectrofluorometrically in real-time (rather than as an endpoint (29)). In addition, conducting the enzyme assay in liquid medium significantly increased scalability while drastically reduced reaction volumes and the amount of substrate and inhibitor needed. In fact, by performing the assay in 96-well plates the amount of substrate and inhibitor used was reduced by 100-fold compared to previous assay methodology (29). While we showcased scalability by conducting the assay in a 96-well format, the assay can be additionally downscaled to suit a 384-well plate, which would further reduce the amount of substrate and inhibitor required (1000-fold reduction over the previous method (29)). The addition of a bacterial growth analysis step (prior to measuring AssT activity and during bacterial treatment with inhibitors) is also easily scalable to fit the 384-well format and would benefit future fragment-based drug design approaches.

In conclusion, our study describes the establishment of a microbiological assay pipeline that can support DsbA inhibitor development all the way from screening to early preclinical candidate validation. Our platform of assays is also suited to screening and evaluating other antivirulence inhibitors (e.g. for flagella components, motility regulators, sulfotransferase activity), while also assessing their potential antibacterial activity (growth inhibition) at the same time. Importantly, we hope our study will serve as a paradigm for the development of similarly accurate, easy to perform, and high throughput cell-based assays that can advance the discovery and preclinical development of other antivirulence drugs that could offer future solutions to curbing the AMR crisis.

## Methods

### Bacterial strains, plasmids, and culture conditions

All bacterial strains utilised in this study (Table 2) were routinely cultured at 37 °C in liquid or on solid lysogeny broth (LB-Lennox) medium supplemented, when required, with chloramphenicol (34 μg/mL) or ampicillin (100 μg/mL), or both. The expression of AssT and DsbA from plasmids pAssT and pEcDsbA, respectively, did not require induction with isopropyl β-D-1-thiogalactopyranoside (IPTG) as the basal level of expression was sufficient for observed effects (27, 28, 29). CFT073 mutants were constructed previously using λ-red-mediated homologous recombination as described elsewhere (28, 49). Plasmids pAssT (27), pEcDsbA (28), and pSU2718 were routinely transformed into strains using electroporation.

**Table 2:** Table of bacterial strains and plasmids.

| *E. coli* strain or plasmid | Description | Reference |
| --- | --- | --- |
| **Strain name** |  |  |
| JCB816 | MC1000 *phoR* λ102 | (50) |
| CFT073 | UPEC isolate (O6:K2:H1) | (51) |
| CFT073Δ*dsbA*Δ*dsbLI* | CFT073Δ*dsbA*::FRTΔ*dsbLI*::FRT | (28) |
| CFT073/pAssT |  | This study |
| CFT073/pSU2718 |  | This study |
| CFT073Δ*dsbA*Δ*dsbLI*/pAssT |  | This study |
| CFT073Δ*dsbA*Δ*dsbLI*/pAssT/pEcDsbA |  | This Study |
| **Plasmids** |  |  |
| pAssT | *assT* gene in pSU2718; Cm<sup>r</sup> | (27) |
| pEcDsbA | *dsbA* gene in pUC19; Ap<sup>r</sup> | (28) |

### Chemicals and stock solutions

Chloramphenicol, ampicillin, phenol, and MUS were purchased from Sigma-Aldrich (Australia), and F1 was purchased from Thermo Fisher Scientific (Australia). MUS (10 mM) and phenol (50 mM) solutions were prepared in sodium chloride (0.9%), and F1 (250 mM) solution was prepared in dimethyl sulfoxide (DMSO). All stock solutions were stored in the absence of light at −20 °C. Working solutions were prepared in LB-Lennox and were used on the same day.

### Absorbance-based bacterial motility assay

Bacterial strains were grown by static 24-hour culture in LB-Lennox media at 37 °C. Cultures were normalized to an O.D. 600 nm of 2 (~2 × 10^9^ CFU/mL) using a spectrophotometer. The multi-well soft agar plates were prepared by adding a volume of 700 μL (24-well) or 350 μL (48-well) of soft LB-Lennox agar (0.25 % [wt/vol]), containing either DMSO (0.4%, vehicle control) or the inhibitor F1 at various concentrations (1-0.1 mM), to each well of the plate. The soft agar was allowed to solidify for at least 2 hours at room temperature (21 °C), before being inoculated in the left-hand corner of each well with 1 μL of normalized culture (~2 × 10^9^ CFU/mL). Inoculated plates were incubated at room temperature for 20 minutes to allow the inoculum to dry. The zone of motility was measured by incubating plates at 37 °C in a CLARIOstar^®^ plate reader (BMG, Australia) programmed to measure absorbance (O.D. 600 nm) at each hour over 15 hours (Figure 5). Absorbance measurements were made using the inbuilt spiral averaging function with orbital averaging producing similar results (data not shown). Instrument data were normalized (with DMSO vehicle control data set at 100%) and plotted using GraphPad Prism 8. Mean motility values were calculated from 3 biological replicates of each strain tested under each specific condition. The start and end of motility were estimated from motility curves and were defined as the beginning and endpoints, respectively, of the exponential phase (zone of motility). The slope of each zone of motility was calculated in Excel and group means were compared for statistical differences by one-way ANOVA (*p* < 0.05) in GraphPad Prism 8. F1 dose-response curves were generated using the motility slopes, and the corresponding IC_50_ value was calculated by applying a non-liner regression (curve fit).

**Figure 5.**
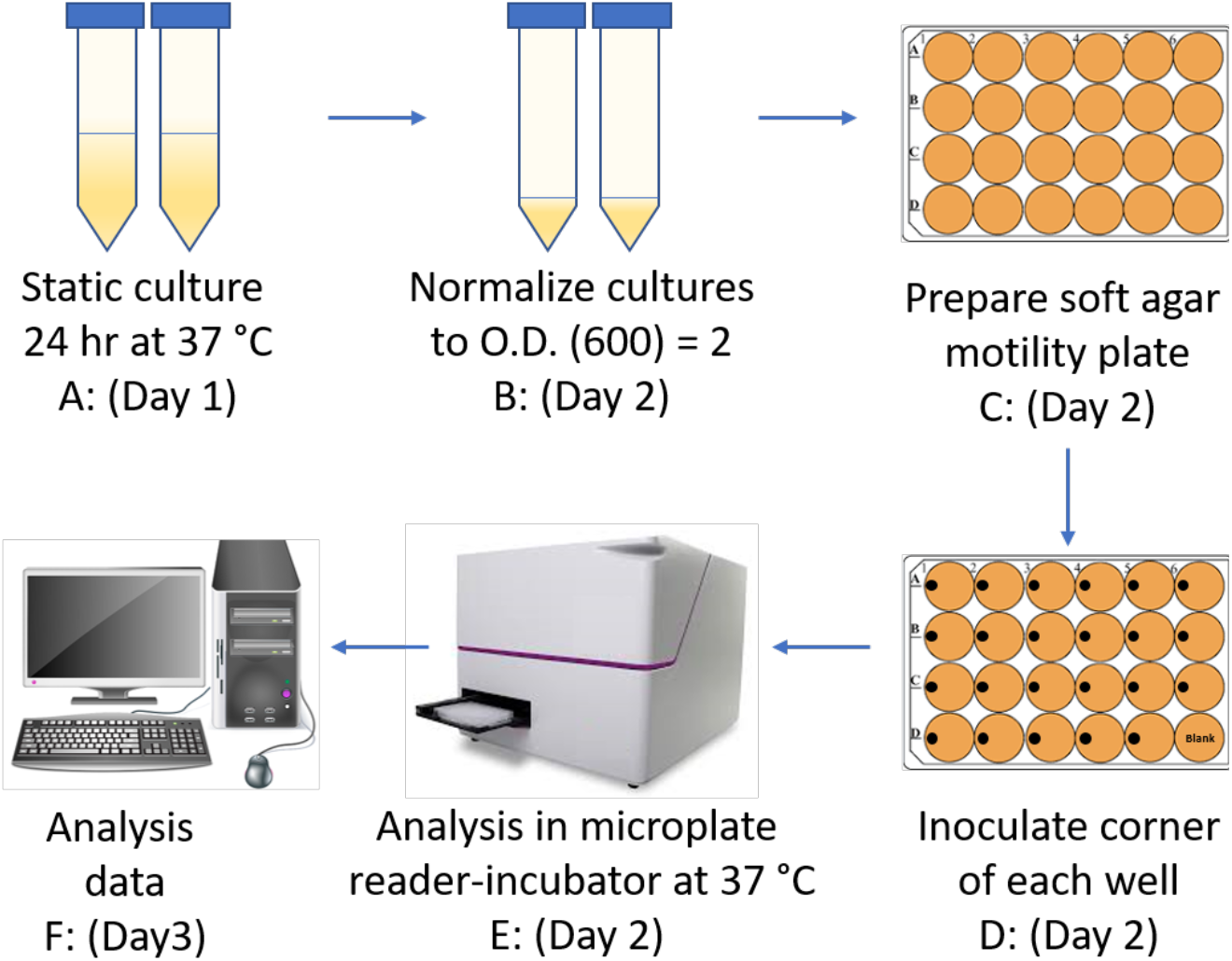
Overivew of absorbance-based bacterial motility assay. (A): Bacterial strains were cultured statically in LB-Lennox media for 24 hours. (B) Overnight cultures were normalized to an O.D. 600 nm of 2. (C) 24-well motility plates were prepared by pipetting 700 μL of warm (55 °C) soft LB agar (0.25%) in each well supplemented with F1 inhibitor or vehicle control (DMSO). The soft agar was allowed to solidify at room temperatures for at least 2 hours. (D) 1 μL of bacterial culture fixed at O.D. 600 nm = 2 was inoculated onto the surface of triplicate soft agar wells by depositing the inoculum at the left edge of the well (with care not to penetrate the agar). The inoculum was allowed to dry onto the agar for 20 minutes at room temperature before the plate was covered with a plastic lid. (E) Bacterial swimming motility was monitored spectrophotometrically for 15-hours in a BMG plate reader at 37°C (using orbital, spiral, or matrix averaging to ensure optimal well coverage). (F) Data acquisition and analysis.

### Cell-based AssT sulfotransferase enzyme assay

Bacterial strains were cultured in LB media, supplemented with antibiotics as appropriate, at 37 °C overnight with shaking at 200 rpm. Overnight cultures were used as inocula in bacterial growth assays (step 1) conducted in a 96-well plate by preparing two-fold serial dilutions of F1 inhibitor compound at twice the desired final concentration in LB-Lennox medium (100 μL final volume). Each well was then inoculated with 100 μL of 1 × 10^7^ CFU/mL inoculum, to give a total well volume of 200 μL and a final cell concentration of 5 × 10^5^ CFU/mL. The growth analysis plate was covered with a breathable sealing membrane (Breathe-Easy® sealing membrane, Sigma, Australia), and incubated at 37 °C for 15 hours with shaking (300 rpm) in a CLARIOstar^®^ plate reader (BMG, Australia) programmed to obtained O.D. 600 nm measurements every 15 minutes over the 15-hour period. At the end of the culture period, each well was normalized to an O.D. 600 nm of 0.4 (~3.5 × 10^8^ CFU/mL) in a fresh 96-well plate (step 2), to ensure that each well contained an equal number of cells. Wells were then supplemented with 4-methylumbelliferyl sulfate (MUS, Sigma, Castle Hill, Australia) (0.5 mM final concentration), and phenol (Sigma, Castle Hill, Australia) (1 mM final concentration) and sulfotransferase activity was monitored immediately in a CLARIOstar^®^ plate reader (BMG, Australia) by measuring fluorescence emitted at 450-480 nm (excitation wavelength at 360-380 nm) and measurements acquired every 5 minutes over a 60 - 90 minute time period (Figure 6). Instrument data were normalized (with DMSO vehicle control set at 100%) and analysed using GraphPad Prism 8. The F1 dose-response curve was generated using fluorescence data from the 40-minute time point, and the corresponding IC_50_ value was calculated by applying a non-linear regression (curve fit).

**Figure 6.**
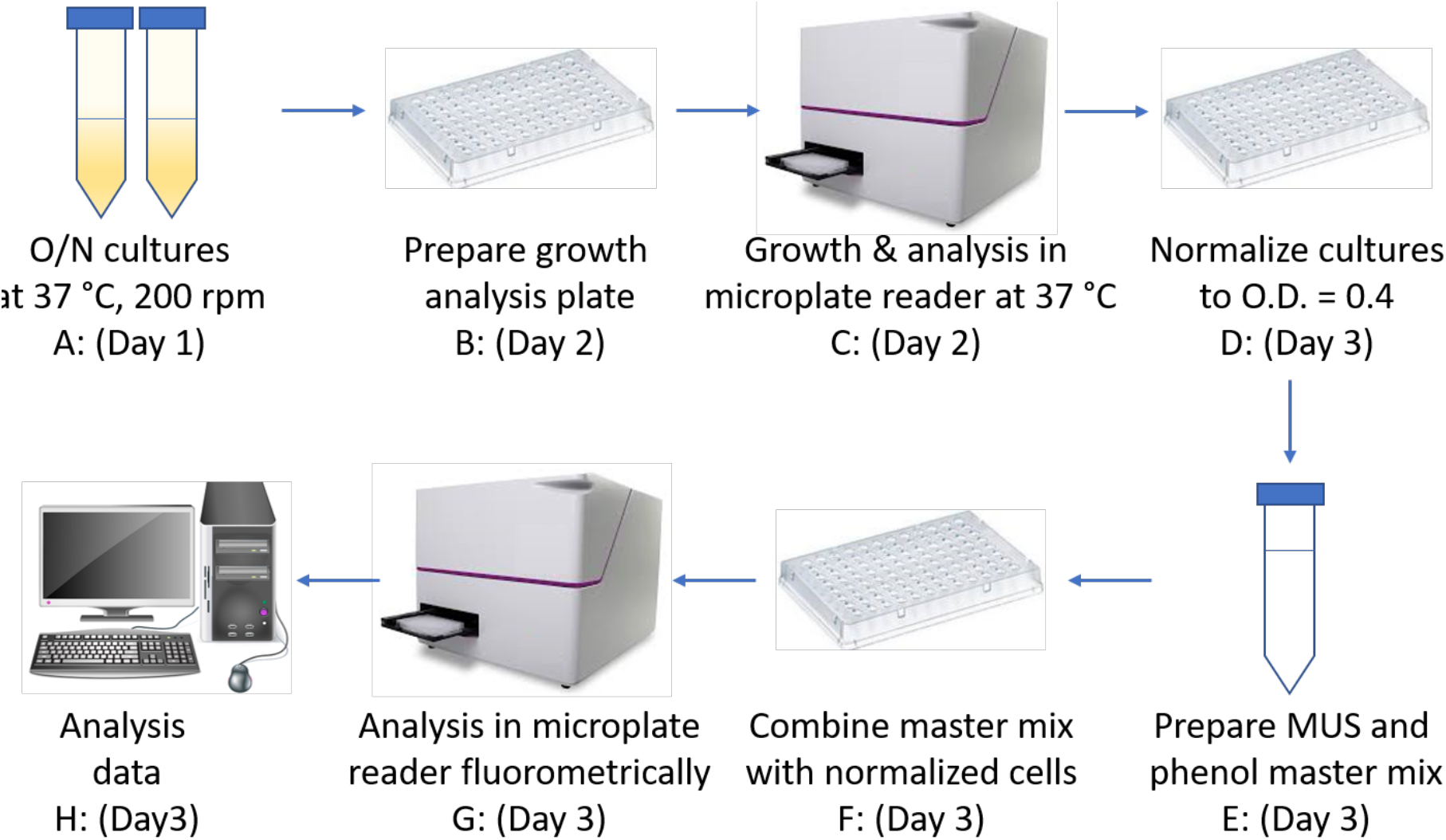
Overview of cell-based AssT sulfotransferase assay. (A) Bacterial strains were cultured in LB-Lennox media overnight at 37 °C with aeration (200 rpm). (B) Growth analysis plates were prepared by subculturing the O/N cultures from (A) into a 96-well plate containing the test inhibitors (akin to preparing an MIC challenge plate). (C) Growth plates were incubated at 37 °C, 300 rpm, in a microplate reader programmed to take O.D. 600 nm readings every 15 minutes for 15 hours. (D) Growth plate cultures were transferred in a fresh 96-well plate with each culture well normalised at an O.D. 600 nm of 0.4. (E) A master mix containing 4 μL of phenol (50 mM), 10 μL of MUS (10 mM), and 126 μL of LB-Lennox per reaction well was prepared. (F) The reaction master mix (140 μL) was added to normalized cultures (60 μL) and mixed. (G) Fluorescence at 450-480 nm was immediately monitored in a CLARIOstar^®^ plate reader (BMG, Australia) with measurements obtained every 5 minutes for up to 90 minutes. (H): Data acquisition and analysis.

## Acknowledgments

This work was supported by a National Health and Medical Research Council Project Grant (APP1144046) and a Clive and Vera Ramaciotti Health Investment Grant (2017HIG0119). MT was supported by a Queensland University of Technology Vice-Chancellor’s Research Fellowship.

